# IL-27 neutralization promotes pro-inflammatory macrophage polarization and improve response to immune checkpoint blockade

**DOI:** 10.1101/2024.09.13.612803

**Authors:** Quentin Glaziou, Laetitia Basset, Sènan d’Almeida, Pascale Pignon, Christophe Blanquart, Yves Delneste, Julie Tabiasco

## Abstract

In this study, we investigated the role of interleukin-27 (IL-27) as a regulator of the tumor immune microenvironment (TME) and its impact on the CD39/adenosine metabolic axis.

The effects of IL-27 neutralization were evaluated using both in vivo MC38 murine colon adenocarcinoma mode and in vitro human macrophage models. In vivo, we analyzed how IL-27 blockade affects tumor growth and the phenotype of infiltrating immune cells, particularly regarding their CD39 expression. In vitro, we focused on the impact of IL-27 during human macrophage differentiation using flow cytometry, metabolic analysis, and functional assays.

Our findings demonstrate that within tumor IL-27 is a major driver of the immunosuppressive phenotype in both murine and human immune cells. Mechanistically, IL-27 induces high levels of the ectonucleotidase CD39, promoting an adenosine-mediated suppressive environment. In vivo, IL-27 blockade led to a broad downregulation of CD39 on both myeloid cells and exhausted T cells. This remodeling of the TME significantly attenuated tumor growth and enhanced the efficacy of anti-PD-L1 checkpoint therapy. We confirm that neutralizing IL-27 prevents the acquisition of immunoregulatory markers in human macrophages and induces significant metabolic reprogramming.

Collectively, our results highlight the IL-27/CD39 axis as a key mechanism of immune evasion. We suggest that targeting IL-27 is a promising strategy to reprogram the metabolic and cellular state of the tumor microenvironment, improving the effectiveness of cancer immunotherapy.

## Introduction

Interleukin-27 (IL-27) has emerged as a paradoxical regulator of anti-tumor immunity. This heterodimeric cytokine, composed of the p28 and EBI3 subunits, belongs to the IL-6/IL-12 cytokine family and signals through a receptor complex formed by IL-27RA and gp130. IL-27 was initially described as a cytokine capable of promoting type 1 immune responses, but subsequent studies demonstrated that it can also induce potent immunoregulatory programs. This dual biology is now considered a defining feature of IL-27: depending on the cellular target, tissue compartment, inflammatory context, timing, and intensity of exposure, IL-27 may either support effector immunity or limit immune activation (1,2).

This apparent contradiction is particularly relevant in cancer. Several studies have shown that IL-27 can promote anti-tumor immunity by enhancing CD8⁺ T cell survival, cytotoxic differentiation, and memory-like features (3,4). More recently, Bréart et al. demonstrated that IL-27 expression is associated with cytotoxic T lymphocyte signatures in human and murine tumors, that IL-27 directly supports tumor-specific CD8⁺ T cells, and that IL-27 receptor agonism using inducible IL-27 expression or a half-life-extended IL-27 protein induces tumor regression and synergizes with PD-L1 blockade (5). In line with this therapeutic concept, systemic IL-27 gene delivery was previously shown to induce anti-tumor activity, promote regulatory T cell depletion, and enhance cancer immunotherapy efficacy (6).

Together, these studies support the idea that strong systemic or therapeutic IL-27 exposure can favor anti-tumor immunity, particularly when it efficiently programs cytotoxic CD8⁺ T cells.

However, IL-27 biology within the tumor microenvironment may not be equivalent to systemic IL-27 agonism. A growing body of evidence indicates that IL-27 can also promote immunoregulatory circuits, including IL-10 production, co-inhibitory receptor expression, PD-L1 expression, and ENTPD1/CD39 upregulation (7–10). In dendritic cells, IL-27 induces ENTPD1/CD39 and suppresses T cell responses (11). In regulatory T cells, IL-27-driven CD39 expression confers protumorigenic activity (12). In macrophages, IL-27 contributes to the acquisition of an immunoregulatory phenotype and promotes CD39 expression during M-CSF-driven differentiation (13). These findings suggest that, within an established and suppressive tumor niche, endogenous IL-27 may participate in a local negative feedback program rather than acting primarily as a CTL-supportive cytokine. This notion is further supported by studies in viral infections, such as influenza, where the timing of IL-27 administration dramatically alters its functional outcome while early IL-27 treatment impairs viral clearance and exacerbates disease, its late administration alleviates immunopathology by dampening the excessive immune response notably reducing neutrophil and monocyte infiltration and inflammatory cytokine production (14).

We therefore reasoned that the apparent duality of IL-27 in cancer may be explained, at least in part, by differences in anatomical compartment, dose, kinetics, and mode of intervention. Systemic IL-27 signaling can influence lymphoid organs, circulating immune cells, tumor-draining lymph nodes, and tumor-infiltrating CTLs (5). In contrast, local IL-27 production within an established tumor microenvironment may preferentially target tumor-associated macrophages, dendritic cells, regulatory T cells, and exhausted T cells, thereby reinforcing immunosuppressive feedback loops as observed in spondyloarthritis or influenza (14,15).

Thus, the biological outcome of IL-27 signaling is likely not determined by the cytokine alone, but by where, when, how strongly, and on which cellular compartment the signal is delivered.

One major candidate mechanism linking local IL-27 signaling to tumor immune suppression is the CD39/CD73/adenosine pathway. Extracellular ATP can act as a pro-inflammatory danger signal, but its sequential degradation by CD39 and CD73 generates adenosine, a potent immunosuppressive metabolite in the tumor microenvironment (16–18). This pathway is active in multiple tumor-associated cell types, including macrophages, myeloid-derived suppressor cells, regulatory T cells, exhausted CD8⁺ T cells, stromal cells, and tumor cells. CD39 is not only an enzymatic driver of adenosine production, but also a marker of highly suppressive Tregs and terminally exhausted tumor-infiltrating CD8⁺ T cells (19–21). Therefore, upstream signals that sustain CD39 expression may coordinate metabolic suppression across several immune compartments simultaneously.

Based on these observations, we hypothesized that endogenous IL-27 locally produced in the tumor microenvironment sustains a CD39-centered suppressive network, distinct from the CTL-supportive effects observed after systemic or therapeutic IL-27 agonism. To test this compartmental hypothesis, we selectively neutralized IL-27 within established MC38 tumors. Rather than assessing systemic IL-27 depletion or therapeutic IL-27 agonism, our study was specifically designed to investigate the local, endogenous role of IL-27 within the tumor microenvironment, with a particular focus on its immunological implications. We confirm that intratumorally IL-27 blockade delays tumor growth in our MC38 model and prolongs survival (22). We show that intratumorally IL-27 blockade enhances the efficacy of PD-L1 blockade. Mechanistically, IL-27 neutralization reduces CD39 expression across tumor-infiltrating myeloid cells and T cells, consistent with attenuation of adenosine-dependent suppression, T cell exhaustion, and Treg-associated suppressive activity. To extend these findings to human cells, we further demonstrate that IL-27 neutralization during primary human macrophage differentiation downregulates CD39/CD73 and other immunosuppressive mediators while promoting a phenotype more permissive to T cell activation and cytotoxicity. These results identify local IL-27 as a regulator of macrophage-driven immune suppression and suggest that intratumorally IL-27 blockade may complement immune checkpoint inhibition by disrupting a CD39-centered suppressive network in the tumor microenvironment.

## Materials and Methods

### Cell Culture

CD14⁺ monocytes and CD4⁺/CD8⁺ T cells were isolated from healthy donor peripheral blood mononuclear cells by positive selection (Miltenyi Biotec). Monocytes were differentiated into macrophages by culturing them for 7 days in RPMI 1640 with 10% fetal bovine serum and 50 ng/mL macrophage colony-stimulating factor (M-CSF) (ORF Genetics, Iceland), a standard concentration for human macrophage generation (23–26). In parallel, a subset of monocytes was differentiated with 50 ng/mL granulocyte–macrophage colony-stimulating factor (GM-CSF) to generate pro-inflammatory macrophages for comparison. IL-27-neutralizing antibody (5 µg/mL; R&D Systems, AF-2526, goat IgG) or isotype control antibody (5 µg/mL; R&D Systems, AB-108-C, goat IgG) was added to monocyte cultures on the day of plating and on day 3 to generate IL-27-neutralized macrophages (Mα27) and isotype control-treated macrophages (Miso), respectively. Where indicated, caffeine (Sigma-Aldrich; 10 µM) or the EP2 receptor antagonist AH6809 (Cayman Chemical; 10 µM) was added during differentiation at the specified concentrations.

### Cell Activation Assays

For macrophage activation, differentiated macrophages were plated at 2 × 10^5^ cells per well in flat-bottom 96-well plates and stimulated with lipopolysaccharide (LPS, E. coli O111:B4; 200 ng/mL; Sigma-Aldrich). Immediately before LPS stimulation, macrophages were extensively washed (3× with pre-warmed PBS [Ca²⁺/Mg²⁺-free] and once with complete medium) to remove any residual neutralizing or isotype antibodies from the differentiation phase. Purified CD4⁺ and CD8⁺ T lymphocytes were seeded in 96-well plates at a density of 2 × 10⁵ cells per well and stimulated with phytohemagglutinin (PHA, 10 µg/mL; Sigma-Aldrich) in antibody-free medium. Stimulation was performed either in the absence or presence of 10% macrophage-conditioned medium (M-CM). M-CM corresponded to supernatants collected after 24 h of LPS activation of Mα27 or Miso macrophages, prepared under antibody-free conditions as described above. Culture supernatants were collected after 48 h and stored at −80 °C prior to cytokine analysis by ELISA.

For T cell proliferation assays, CD4⁺ T cells were labeled with 3 µM CFDA-SE (Invitrogen) and cultured (2 × 10^5^ cells per well) in 96-well plates pre-coated with anti-CD3 antibody (1 µg/mL, clone OKT3; ATCC). Soluble anti-CD28 (1 µg/mL; BD Pharmingen) was added to provide co-stimulation. Macrophage influence was tested by adding 50% (v/v) of M-CM from 24 h LPS-activated Mα27, Miso, or macrophage without antibody addition (untreated macrophages) cultures (all M-CM generated after the extensive washing step and in antibody-free medium). After 6 days, CD4⁺ T cell proliferation was assessed by flow cytometry based on CFDA-SE dilution.

For CD8⁺ T cell cytotoxicity assays, a luciferase-expressing MUC1⁺ murine mesothelioma cell line (Meso34) was co-cultured with a MUC1-specific CD8⁺ T cell clone (26) (1 × 10^5^ T cells with 3 × 10^4^ target cells) in the presence or absence of macrophages (2 × 10^4^ Mα27 or Miso). Before co-culture, macrophages were thoroughly washed as above and resuspended in antibody-free assay medium to preclude any antibody carryover. Co-cultures were maintained for 72 h. Tumor cell killing was quantified by measuring luciferase activity released into the supernatant, which correlates with tumor cell lysis.

Across all functional assays, “Miso” macrophages served as isotype-exposed controls during differentiation, and no anti–IL-27 or isotype IgG was present during LPS activation, in the conditioned media, or during T cell co-culture. These procedures ensure that differences between Mα27 and Miso reflect macrophage-derived factors and phenotype rather than direct antibody effects.

### Soluble Mediator Analysis

Concentrations of cytokines and other mediators in culture supernatants were measured by ELISA according to the manufacturers’ protocols. The following kits were used: human IL-10, IL-2, IFN-γ (Diaclone); human CCL2, CCL22, prostaglandin E₂ (PGE₂), IL-27 (R&D Systems). For the quantification of membrane-associated IL-27, cell pellets were subjected to five freeze–thaw cycles, resuspended in PBS, disrupted by sonication, and centrifuged at 10^5^ × g for 10 min. The clarified supernatants were subsequently used for ELISA analysis.

### Western Blotting

Macrophage cell lysates (1 × 10^6^ cells per sample) were prepared in RIPA lysis buffer with protease/phosphatase inhibitors. Equal amounts of protein were separated by SDS-PAGE and transferred to PVDF membranes. Blots were probed with primary antibodies against human COX-2 and CD73 (both 1 µg/mL; Cell Signaling Technology), followed by HRP-conjugated secondary antibodies (anti-rabbit IgG or anti-mouse IgG, 1 µg/mL; Cell Signaling). β-actin (1 µg/mL; Cell Signaling) was used as a loading control. Bands were visualized using ECL or SuperSignal West Femto chemiluminescent substrate (Thermo Fisher).

### Gene Expression Analysis

Total RNA was extracted using the RNeasy Mini kit (Qiagen) and reverse-transcribed with Superscript II (Thermo Fisher). Quantitative PCR was performed using SYBR Green Supermix (Bio-Rad) on a CFX96 thermocycler (Bio-Rad). Primer sequences (e.g., for IL10, NT5E/CD73, PTGS2/COX-2) are available on request. Gene expression was calculated by the 2^(-ΔCt)^ method, normalizing to GAPDH as a housekeeping gene.

### Metabolic Assays

Macrophage metabolic function was assessed using a Seahorse XFe24 Analyzer (Agilent). Oxygen consumption rate (OCR) was measured with the Seahorse Mito Stress Test, and extracellular acidification rate (ECAR) was measured with the Glycolysis Stress Test, following the manufacturer’s protocols. Macrophages were plated at 1 × 10^5^ cells per well in Seahorse plates and sequentially treated with oligomycin, FCCP, and rotenone/antimycin A (for OCR assays) or glucose, oligomycin, and 2-deoxyglucose (for ECAR assays). Parameters of mitochondrial respiration (basal OCR, maximal OCR, ATP-linked OCR) and glycolysis (basal ECAR, glycolytic capacity) were derived. Lactate production was measured in 24 h culture supernatants (from 1 × 10^5^ cells per well) using a colorimetric L-lactate assay kit (Abcam).

### In Vivo Tumor Model

The MC38 murine colon adenocarcinoma cell line was maintained in culture and harvested during log-phase for injection. Female C57BL/6 mice (8 weeks old) were injected subcutaneously in the right flank with 5 × 10^5^ MC38 cells in 100 µL PBS. Once tumors became detectable in all mice (approximately 8– 10 days post-injection), animals were randomly divided into two groups. One group received an IL-27-neutralizing antibody (anti-mouse IL-27p28, clone MM27-7B1, Ultra-LEAF purified; BioLegend), while the other received a non-relevant isotype control antibody of the same IgG subclass (Mouse IgG2a, κ Isotype Control, Clone: MOPC-173, Ultra LEAF purified; Biolegend). Treatments were administered intratumorally every 48 hours for a total of five doses, each consisting of 20 µg in 20 µL per injection. Starting two days after the final anti-IL-27 injection, an anti-PD-L1 antibody (clone 10F.9G2, InVivoMab; BioXcell) was given intraperitoneally (200 µg in 200 µL per injection) every three days for three doses. Tumor growth was monitored every two days using calipers. Tumor volume was calculated as (length × width × height) × 0.52 (“0.52” comes from π/6 for an ellipsoid approximation). Tumor measurements were performed by two independent researchers, one of whom was blinded to the treatment groups. Mice were euthanized by cervical dislocation when tumor volume exceeded 1,800 mm³ or if tumors ulcerated.

### Flow Cytometry Human macrophages

Macrophages were stained with fluorochrome-conjugated antibodies (Supplementary Table 1). Non-viable events were excluded with 7-AAD (BioLegend, San Diego, CA, USA). Acquisition was performed on a BD FACSCanto flow cytometer (BD Biosciences, San Jose, CA, USA). Data were analyzed in FlowJo v10.9 (FlowJo LLC). Median fluorescence intensity (MFI) values for each marker were normalized to the MFI of the corresponding isotype control on the same sample, yielding a relative fluorescence intensity (RFI). These RFI values were used to compare expression between Mα27 and Miso conditions.

### Tumor-infiltrating cells

Tumors were minced with scissors and enzymatically dissociated (collagenase IV + DNase I, 20 min, 37 °C). Cells were washed and passed through 200 µm strainers, then stained with LIVE/DEAD Fixable Yellow (Thermo Fisher Scientific) to exclude non-viable events. Fc receptors were blocked (FcR Block, Miltenyi Biotec, Germany) before surface staining with the antibody panels listed in Supplementary Table 1. Multiparameter flow cytometry was performed on a BD LSRFortessa (BD Biosciences). Median fluorescence intensities (MFIs) were calculated on viable cells; data were analyzed in FlowJo v10 (FlowJo LLC).

### Statistics

Statistical analyses were performed in GraphPad Prism 10 (GraphPad Software, San Diego, CA, USA). Two-group comparisons used the Mann–Whitney U test. For comparisons involving three or more groups, one-way ANOVA with Tukey’s multiple-comparisons test was applied. Survival curves were compared using the log-rank (Mantel–Cox) test. All tests were two-sided, and statistical significance was set at α = 0.05 (p < 0.05; *p < 0.01; **p < 0.001). In the graphical representations, only statistically significant differences directly relevant to the discussion were displayed. Results not reaching statistical significance but showing a consistent pattern were reported in the text and described as trends.

## Results

### In vivo IL-27 neutralization delays tumor growth and prolongs survival

Mice bearing established MC38 tumors were treated intratumorally with an IL-27-neutralizing antibody or control IgG (Figure 1A). IL-27 neutralization resulted in significantly slower tumor growth compared the control treatment (Figure 1B, 1C). By day 10, IL-27-neutralizing antibody treatment significantly delayed tumor growth relative to mice receiving an isotype control (Fig. 1B–C). Tumor growth curves diverged as early as 6 days after starting treatment, indicating a relatively rapid effect of IL-27 neutralization (Figure 1D). Furthermore, IL-27 antibody-treated mice showed a survival benefit. The median survival in the anti-IL-27 group was 18.5 days, compared to 14 days in controls (Figure 1E). This extension of survival was statistically significant (log-rank *p* = 0.008). Thus, neutralizing IL-27 in the TME can impede tumor progression and modestly improve survival even as a single agent.

**Figure 1.**
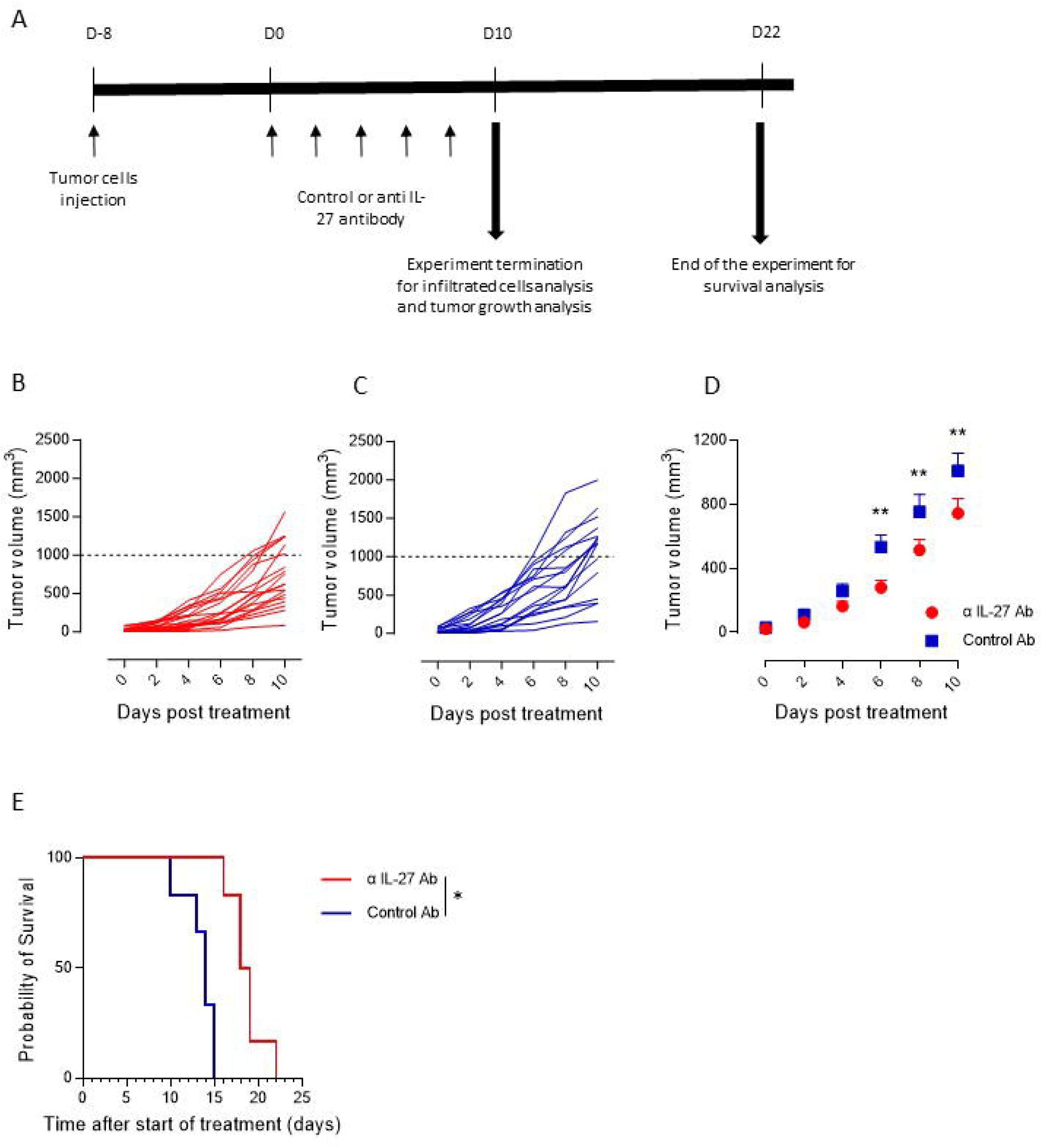
Therapeutic efficacy of anti-IL-27 in MC38 tumors. (A) 5 × 10^5^ MC38 cells injected s.c. in the right flank of C57BL/6 mice. Eight days later (D0), mice received intratumoral αIL-27 or isotype-matched control IgG every 48 h (D0, D2, D4, D6, D8; five doses total). Tumor monitoring ended when the first tumor reached 1,800 mm³ (D10); survival follow-up ended at D22 when the last tumor reached 1,800 mm^3^. (B–C) Individual tumor-growth spaghetti plots by treatment arm (αIL-27 vs Control Ab). (D) Cumulative mean tumor volumes (± SEM) every two days (n = 19 for treated group, n = 17 for control group). (E) Kaplan–Meier survival of MC38-bearing mice by treatment. Each symbol = one mouse. Data pooled from two independent experiments; mean ± SEM. Statistics: one-way ANOVA with Tukey’s post-hoc for volumes; log-rank (Mantel–Cox) for survival (overall p = 0.008). Significance: p < 0.05 (*p<0.05, **p<0.01).

### IL-27 blockade modulates the tumor immune microenvironment

To understand how IL-27 neutralization affects immune cells in the tumor, we analyzed immune infiltrates at day 10 of therapy. We focused on CD39 expression, as CD39 produces immunosuppressive adenosine and is regulated by IL-27 and is generally associated with an immunoregulatory phenotype in the cells expressing it.

Overall, tumors treated with anti-IL-27 had lower levels of CD39 on infiltrating myeloid cells. Specifically, CD39 mean fluorescence intensity (MFI) was significantly reduced on newly recruited Ly6C+ F4/80+ macrophages, dendritic cells (DCs), and neutrophils (Figure 2A, 2B, 2C). Mature tumor-associated macrophages (TAMs, CD11b^+^F4/80^high^Ly6C^−^) also showed a trend toward lower CD39 expression (Figure 2D). These results suggest that IL-27 blockade limits the metabolic suppressive capacity of the myeloid compartment.

**Figure 2.**
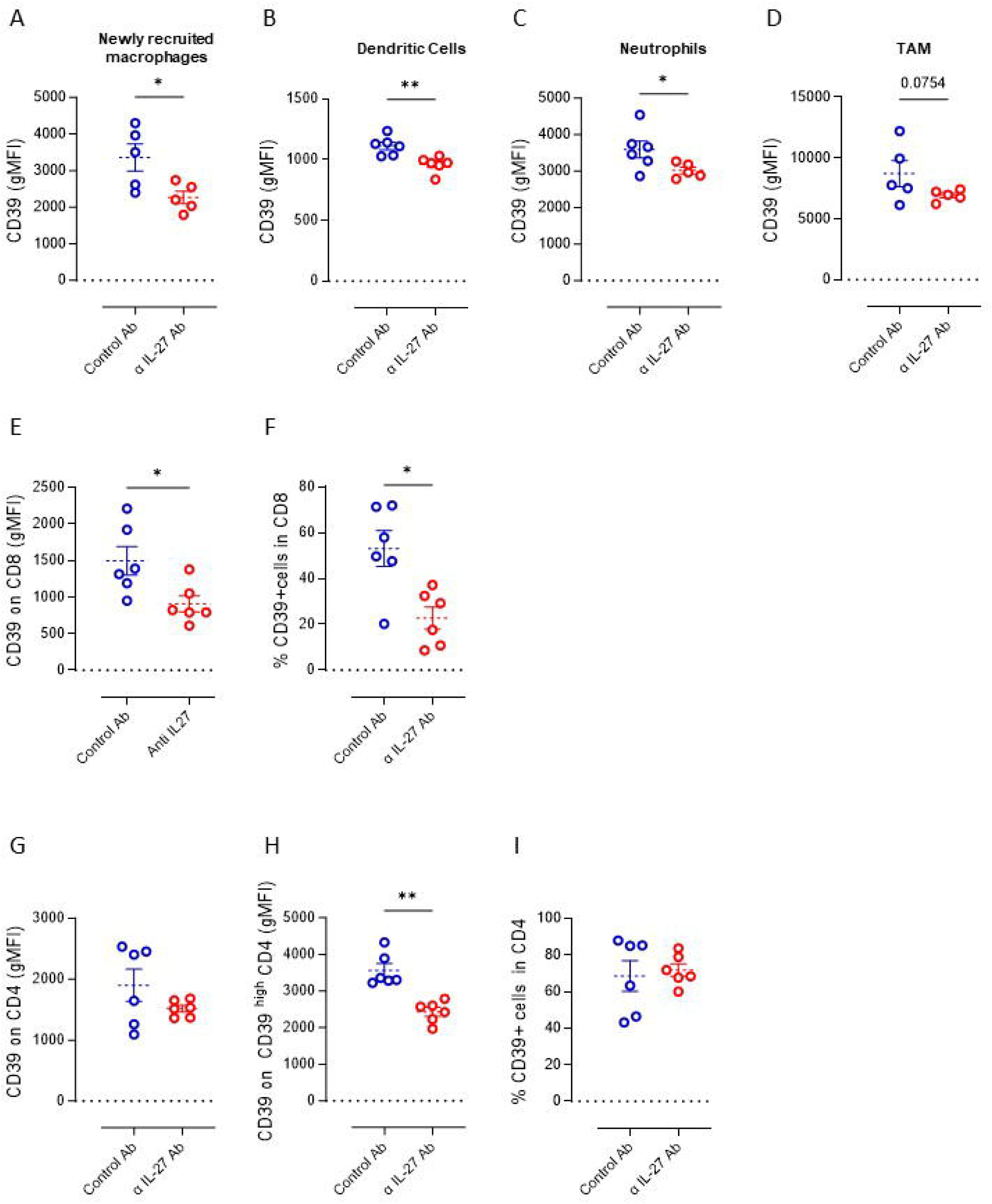
IL-27 neutralization lowers CD39 on tumor-infiltrating immune cells. Upon tumors reaching 20–50 mm³ (∼day 8), mice received intratumoral αIL-27 or isotype control every 48 h (D0, D2, D4, D6, D8). Tumors were harvested on day 10 for flow cytometry. (A–D) CD39 gMFI on neutrophils, dendritic cells, monocytes and TAM. (E) CD39 gMFI on infiltrating CD8⁺ T cells. (F) Frequency of CD39⁺ cells among CD8⁺ T cells. (G) CD39 gMFI on infiltrating CD4⁺ T cells. (H) CD39 gMFI on CD4⁺ T cells within the CD39^high^ subset. (I) Frequency of CD39⁺ cells among CD4⁺ T cells. Each symbol = one mouse. Data pooled from two independent experiments; mean ± SEM. Statistics: Mann–Whitney tests. Significance: p < 0.05 (*p<0.05, **p<0.01).

We also observed notable effects on T cells. CD8^+^ T cells from anti-IL-27-treated tumors showed lower surface levels of CD39 (Figure 2E) and a significant decrease in the percentage of CD39^+^ cells (Figure 2F). In the tumor microenvironment, CD39 is a key marker of terminal exhaustion for CD8^+^ T cells. These CD39^+^ cells usually co-express other inhibitory receptors (like PD-1 or TIGIT) and have poor cytotoxic function. Therefore, the reduction of CD39 suggests that IL-27 neutralization prevents CD8^+^ T cells from reaching a state of terminal exhaustion, potentially keeping them more functional.

Regarding CD4^+^ T cells, while the average level of CD39 did not change for the total population (Figure 2G), the subset with the highest CD39 expression was significantly reduced (Figure 2H). Because CD39 is a classic marker used to identify highly suppressive regulatory T cells (Tregs) in tumors, this reduction suggests a decrease in the suppressive strength of the Treg compartment. The total frequency of CD39^+^ cells among CD4^+^ T cells remained unchanged (Figure 2I), indicating that IL-27 blockade primarily affects the level of CD39 per cell in these suppressive subsets rather than eliminating the cells entirely.

In summary, IL-27 neutralization in vivo induces changes consistent with a less suppressive TME: myeloid cells show lower CD39, and T cells exhibit markers of reduced exhaustion and lower Treg activity.

### IL-27 neutralization enhances the efficacy of PD-L1 blockade

We then tested if neutralizing IL-27 could improve the effect of PD-L1 blockade. A separate group of mice was treated with anti-IL-27, followed by anti-PD-L1 antibody (Figure 3A). The combination of IL-27 neutralization and PD-L1 blockade resulted in stronger tumor growth inhibition than either treatment alone (Figure 3B).

**Figure 3.**
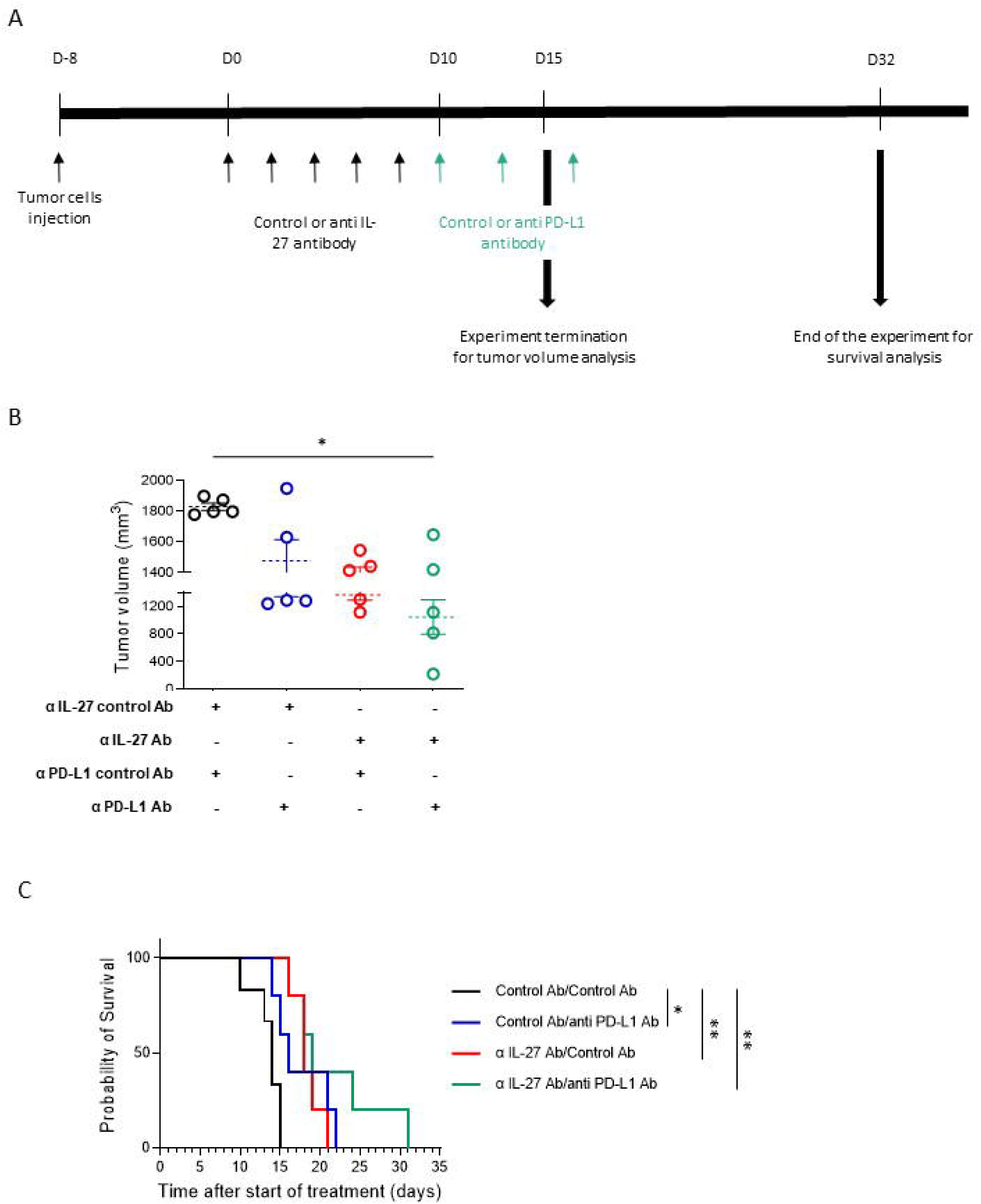
Combination therapy: anti-IL-27 enhances response to anti-PD-L1. (A) MC38 model as in Figure 4. After five intratumoral doses of αIL-27 or isotype control (D0–D8), anti-PD-L1 was started 2 days later (i.p. every 3 days on D9, D12, D15). Tumor monitoring ended when the first tumor reached 1,800 mm³ (D15); survival follow-up ended at D31. (B) Tumor volumes at D15 by treatment group. (C) Kaplan–Meier survival by treatment group. Each symbol = one mouse. Data pooled from two independent experiments; mean ± SEM. Statistics: one-way ANOVA with Tukey’s post-hoc for volumes; log-rank for survival (exact p values annotated on the plot; e.g., *p = 0.04; **p = 0.002). Significance: p < 0.05 (*p<0.05).

At the end of the study, tumors in the combination group were significantly smaller than those in the control group and tended to be smaller than those in mice receiving anti-PD-L1 alone (Figure 3B). While each monotherapy was effective, the combination showed an additional benefit, with survival curves suggesting a trend toward longer survival compared to anti-PD-L1 alone (Figure 3C). These findings suggest that IL-27 neutralization can sensitize the tumor environment to checkpoint inhibition. In summary, IL-27 blockade slows tumor growth and enhances the effect of PD-L1 therapy. These results are consistent with the idea that targeting IL-27 helps reduce immunosuppression in the TME and improves the response to immunotherapy.

### IL-27 neutralization during macrophage differentiation reduces immunosuppressive properties

To translate our in vivo findings, we investigated whether IL-27 neutralization similarly affects human primary cells. While the impact of IL-27 on human CD4+ and CD8+ T cell exhaustion and CD39 expression is already well-documented, its role in human macrophage differentiation remains less understood. Therefore, we focused our human studies on the myeloid compartment to characterize this specific mechanism of immune regulation.

### Characterization of the IL-27 axis in human monocytes

We first confirmed that human monocytes express and release IL-27. RT-qPCR showed the presence of transcripts for both IL-27 subunits (p28 and Ebi3), as well as its receptors (IL−27RA and gp130), in monocytes before differentiation (Supplementary Figure 1A). We observed that IL-27p28 protein concentration increased progressively in the culture supernatants during M-CSF-driven differentiation (Supplementary Figure 1B). While IL-27 was undetectable in supernatants from unstimulated cells at day 5 (Supplementary Figure 1C), we detected IL-27 p28 and Il-27 RA mRNA within the cells at day 5 (Supplementary figure 1D), and found that IL-27p28 was present at the cell surface (Supplementary Figure 1E), suggesting an autocrine or paracrine signaling loop.

### IL-27 blockade downregulates the CD39/CD73 adenosine axis

To assess the impact of IL-27 on macrophage polarization, we differentiated human monocytes with M-CSF (50 ng/mL) and neutralized IL-27 throughout this process. Consistent with our previous findings (13), macrophages differentiated with IL-27 neutralization (Mα27) showed significantly reduced CD39 expression compared to isotype-treated controls (Miso). Flow cytometry confirmed that surface CD39 protein levels were lower on Mα27 cells than on Miso cells, and qRT-PCR analysis similarly revealed a decrease in ENTPD1 (CD39) transcripts in Mα27 (consistent with ref. (13); data not shown). Because adenosine signaling is linked to CD39 function in TAMs, we next tested whether disrupting adenosine signaling would similarly affect CD39 expression. Indeed, adding caffeine, a non-selective adenosine receptor antagonist, during macrophage differentiation significantly decreased surface CD39 levels (Supplementary Figure 2A), supporting the role of adenosine in maintaining CD39^high^ immunosuppressive macrophages (27).

We next examined CD73, the cell-surface ectoenzyme that converts AMP (the product of CD39) into adenosine. IL-27 neutralization led to significantly lower CD73 surface expression on Mα27 macrophages compared to Miso controls (Figure 4A, 4B). This decrease was also evident at the transcript level: qRT-PCR showed that NT5E (CD73) mRNA was reduced in Mα27 cells relative to Miso (Supplementary Figure 3A). Together with our previous report (13), these results indicate that IL-27 signaling promotes high expression of both ectonucleotidases in the adenosine pathway (CD39 and CD73). Consequently, blocking IL-27 during macrophage differentiation downregulated both CD39 and CD73, potentially limiting the cells’ capacity to generate immunosuppressive adenosine.

**Figure 4.**
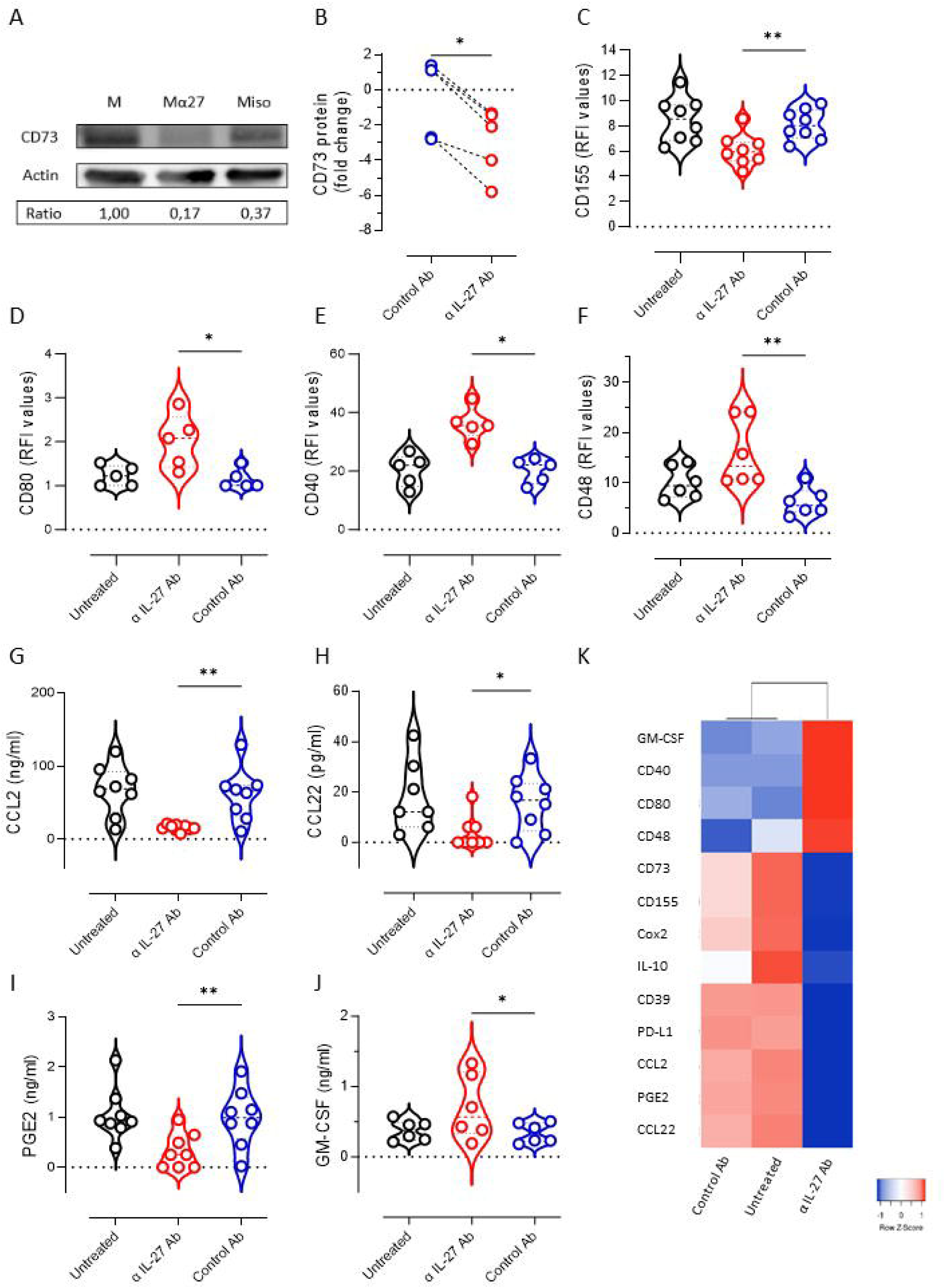
IL-27 neutralization during macrophage differentiation remodels phenotype. Macrophages were differentiated for 7 days with M-CSF alone (Untreated), M-CSF + αIL-27 antibody (αIL-27 Ab), or M-CSF + isotype-matched control IgG (Control Ab). (A) CD73 protein (representative donor) quantified by western blot. (B) Paired comparison of CD73 protein levels relative to Untreated (Wilcoxon matched-pairs signed-rank test). (C–F) Surface expression of immunomodulatory molecules (CD155, CD80, CD40, CD48) by flow cytometry on CD14⁺ gated cells. (G–H) Secreted CCL2 and CCL22 after 24 h culture (ELISA). (I–J) After 24 h LPS stimulation, secreted PGE₂ and GM-CSF (ELISA). (K) Heatmap summarizing significant phenotype changes with IL-27 neutralization. Flow-cytometry data are shown as relative fluorescence intensity (RFI) = MFI_specific / MFI_isotype; qRT-PCR as mRNA normalized to GAPDH; ELISA in ng/mL. Each dot = an independent experiment. Data are mean ± SEM. Statistics: one-way ANOVA with Tukey’s post-hoc; paired Wilcoxon for panel B. Significance: p < 0.05 (*p < 0.05; **p < 0.01; ***p < 0.001).

While we previously showed that IL-27 neutralization reduces PD-L1 expression on human macrophages (13), we now also find that IL-27-neutralized macrophages display diminished CD155 expression (Figure 4C). CD155 (PVR) is an immune checkpoint ligand that interacts with receptors such as TIGIT on T and NK cells, typically delivering an inhibitory signal (28–30). In cancers, high CD155 expression is associated with T cell dysfunction and resistance to immunotherapy (28,31). In our experiments, Mα27 macrophages showed roughly a 25% reduction in CD155 surface levels compared to Miso (Figure 4C).

Conversely, IL-27 neutralization increased the expression of several co-stimulatory or pro-inflammatory surface molecules on macrophages. Mα27 cells had significantly higher levels of CD80 and CD40 than Miso cells (Figure 4D, 4E). CD80 provides a co-stimulatory signal to T cells (via CD28) and is typically associated with classically activated M1 macrophages. CD40, a TNF receptor family member, is an activation-induced molecule on macrophages and dendritic cells that can license these cells to better prime T cells via CD40L interactions (32). We also observed a moderate but significant increase in CD48 on Mα27 macrophages (Figure 4F). CD48 can function as an activating receptor on immune cells and engage CD244 (2B4) on NK/T cells to promote immune activation in certain contexts (33).

In addition to cell surface markers, we analyzed the secretion of key immunoregulatory mediators by macrophages. Without LPS stimulation, Mα27 macrophages secreted significantly less CCL2 than Miso macrophages (Figure 4G). CCL2 is a chemokine that recruits monocytes to tumors, where they can differentiate into TAMs; thus, lower baseline CCL2 in Mα27 suggests a less chemoattractant, pro-tumoral profile. Upon LPS activation, however, Mα27 secreted more CCL2 than Miso (Supplementary Figure 3B), a pattern similar to inflammatory macrophages which produce little CCL2 at rest but upregulate it during activation (Supplementary Figure 3C). Another chemokine, CCL22, which recruits CCR4⁺ regulatory T cells, was found at significantly lower levels in supernatants from Mα27 versus Miso macrophages (Figure 4H). Notably, IL-27 produced by dendritic cells has been shown to induce CCL22 and facilitate Tregs accumulation in tumors (34), suggesting that IL-27-neutralized macrophages might be less effective at attracting Tregs.

We confirmed that IL-27-neutralized macrophages produce less IL-10 upon LPS activation than controls (Supplementary Figure 3D), consistent with our previous report (13). IL-10 is a key immunosuppressive cytokine associated with M2-like macrophages, so its reduction further indicates an attenuated regulatory phenotype in Mα27 cells. On the other hand, we examined two factors that can modulate anti-tumor immunity: prostaglandin E₂ (PGE₂) and GM-CSF.

PGE₂, generally immunosuppressive in the TME, is involved in TAM development (35); inhibiting PGE₂ signaling via EP2 receptor antagonist AH6809 significantly reduced CD39 levels during macrophage differentiation (Supplementary Figure 2B). We found that LPS-stimulated Mα27 macrophages secreted substantially less PGE₂ than Miso (Figure 4I). Correspondingly, expression of cyclooxygenase-2 (COX-2, PTGS2), the enzyme responsible for PGE₂ synthesis, was markedly downregulated in Mα27 at both the mRNA and protein levels (Supplementary Figures 3E, 3F, 3G). This suggests that IL-27 blockade skews macrophages away from a COX-2^high^, PGE₂-producing state. GM-CSF is a cytokine that can drive monocytes toward dendritic cell differentiation and is linked to improved antigen presentation. We observed that Mα27 macrophages secreted significantly more GM-CSF than Miso upon LPS stimulation (Figure 4J). Notably, adding a low dose of GM-CSF during M-CSF-driven differentiation drastically reduced CD39 expression (Supplementary Figure 2C), consistent with GM-CSF’s role in promoting a pro-inflammatory macrophage program (36). The elevated GM-CSF production by Mα27 cells may thus amplify local immune activation (by recruiting/activating dendritic cells, for example).

Collectively, these results demonstrate that IL-27 neutralization during human macrophage differentiation induces a broad phenotypic shift: reduced expression of immunosuppressive molecules (CD39, CD73, CD155, IL-10, CCL22, PGE₂/COX-2) and increased expression of immunostimulatory molecules (CD80, CD40, CD48, GM-CSF, and inducible CCL2). A heatmap summarizing these changes is shown in Figure 4K. Overall, targeting IL-27 yielded macrophages with significantly diminished immunosuppressive properties.

### IL-27 neutralization alters macrophage metabolism toward a pro-inflammatory profile

Cellular metabolism is closely tied to macrophage activation status. Inflammatory activation of murine macrophages typically suppresses oxidative phosphorylation (OxPhos) while enhancing glycolysis, driven by TCA-cycle rewiring (succinate/citrate, itaconate) and nitric oxide–mediated respiratory inhibition. In humans, this split is less stark, glycolysis often rises without loss of substantial OxPhos, highlighting bioenergetic plasticity (37). Metabolic programming is therefore a key determinant of immune function (38–40).

As a baseline comparison, we first characterized metabolic differences between GM-CSF– differentiated macrophages and M-CSF–differentiated macrophages. Consistent with a more activated phenotype, GM-CSF macrophages had significantly higher basal OCR (oxygen consumption in the resting state) and ATP-linked OCR than M-CSF macrophages (Figure 5A, 5B). This indicates that GM-CSF-conditioned macrophages were more metabolically active at rest, engaging oxidative phosphorylation to a greater degree. Interestingly, maximal OCR (mitochondrial respiratory capacity) did not differ between GM-CSF and M-CSF groups (Figure 5C), suggesting GM-CSF macrophages utilize more of their mitochondrial capacity at baseline without necessarily having greater total capacity. We also found that GM-CSF macrophages displayed higher basal ECAR (glycolytic rate) and glycolytic capacity compared to M-CSF macrophages (Figure 5D, 5E). Correspondingly, GM-CSF macrophages produced more lactate in culture (Figure 5F). Thus, GM-CSF-derived macrophages had elevated utilization of both oxidative and glycolytic pathways relative to M-CSF macrophages, aligning with a more pro-inflammatory metabolic profile.

**Figure 5.**
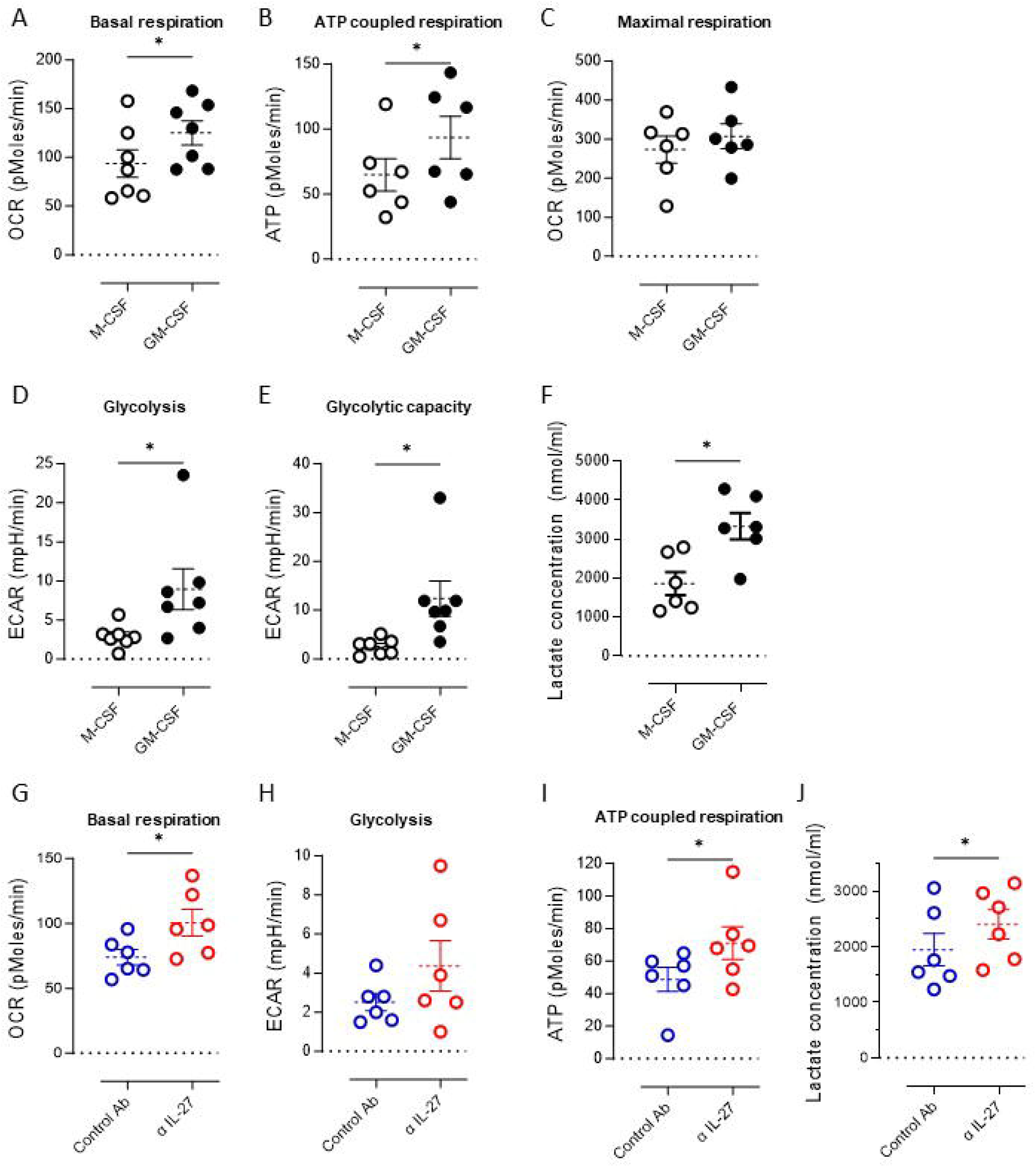
IL-27 neutralization alters macrophage metabolism. (A–C) Oxygen consumption rate (OCR) parameters from Seahorse XF Cell Mito Stress Test, namely basal respiration, maximal respiration, ATP-linked respiration, in macrophages differentiated with M-CSF or GM-CSF. (D–E) Extracellular acidification rate (ECAR) parameters from Glycolysis Stress Test, namely glycolysis and glycolytic capacity, in M-CSF vs GM-CSF conditions. (F) Lactate concentration (supernatants) in M-CSF vs GM-CSF macrophages. (G, I) OCR parameters (basal, ATP-linked) in Miso vs Mα27 macrophages. (H) Basal ECAR in Miso vs Mα27. (J) Lactate concentration in Miso vs Mα27. Each dot = an independent experiment; mean ± SEM. Statistics: unpaired two-tailed Student’s ***t***-test (GraphPad Prism 10) for all two-group comparisons. Significance: p < 0.05 (*p<0.05).

We then assessed the metabolism of IL-27-neutralized versus control antibody M-CSF macrophages. Even though both groups were generated with M-CSF, IL-27 neutralization significantly altered their metabolic state. Mα27 macrophages showed higher basal OCR than Miso, indicating enhanced mitochondrial respiration (Figure 5G). There was also a trend toward increased basal ECAR (glycolysis) in Mα27 (Figure 5H), though this did not reach significance. Importantly, ATP production (estimated from the oligomycin-sensitive OCR) was significantly greater in Mα27 compared to Miso (Figure 5I), as was lactate secretion (Figure 5J). The latter indicates a higher glycolytic flux in Mα27 cells. Together, these data suggest that IL-27-neutralized macrophages are more energetically active, with a metabolism shifted somewhat toward that of pro-inflammatory macrophages. Indeed, the metabolic differences between Mα27 and Miso (higher OCR, higher lactate) mirrored the direction of change seen between GM-CSF and M-CSF macrophages. These results imply that IL-27 blockade drives metabolic reprogramming consistent with macrophage activation and may support the functional changes (e.g., greater cytokine production) observed in Mα27 cells.

### IL-27-neutralized macrophages better support T cell activation and cytotoxicity

The phenotypic and metabolic reprogramming of macrophages by IL-27 neutralization suggested that these cells might be less suppressive toward T cells. First, we examined CD8⁺ T cell effector function in the presence of M-CM. When activated CD8⁺ T cells were cultured with 10% of control M-CM, their secretion of IFN-γ was almost completely inhibited (Supplementary Figure 4A) indicating that factors produced by control macrophages can strongly suppress CD8⁺ T cells. By contrast, under the same conditions, CD8⁺ T cells cultured with supernatant from activated Mα27 macrophages secreted significantly more IFN-γ than those with Miso supernatant (Figure 6A). Although IFN-γ production was still reduced relative to T cells with no macrophage supernatant (Supplementary Figure 4A), the suppression was markedly less severe with Mα27-conditioned medium. This finding suggests that IL-27-neutralized macrophages produce fewer soluble suppressive factors that inhibit CD8⁺ T cell cytokine release. To preclude antibody-related artifacts, macrophages were extensively washed immediately before the 24-h LPS activation used to generate M-CM, thus, the differences observed in T-cell readouts reflect macrophage-derived factors rather than antibody carryover.

**Figure 6.**
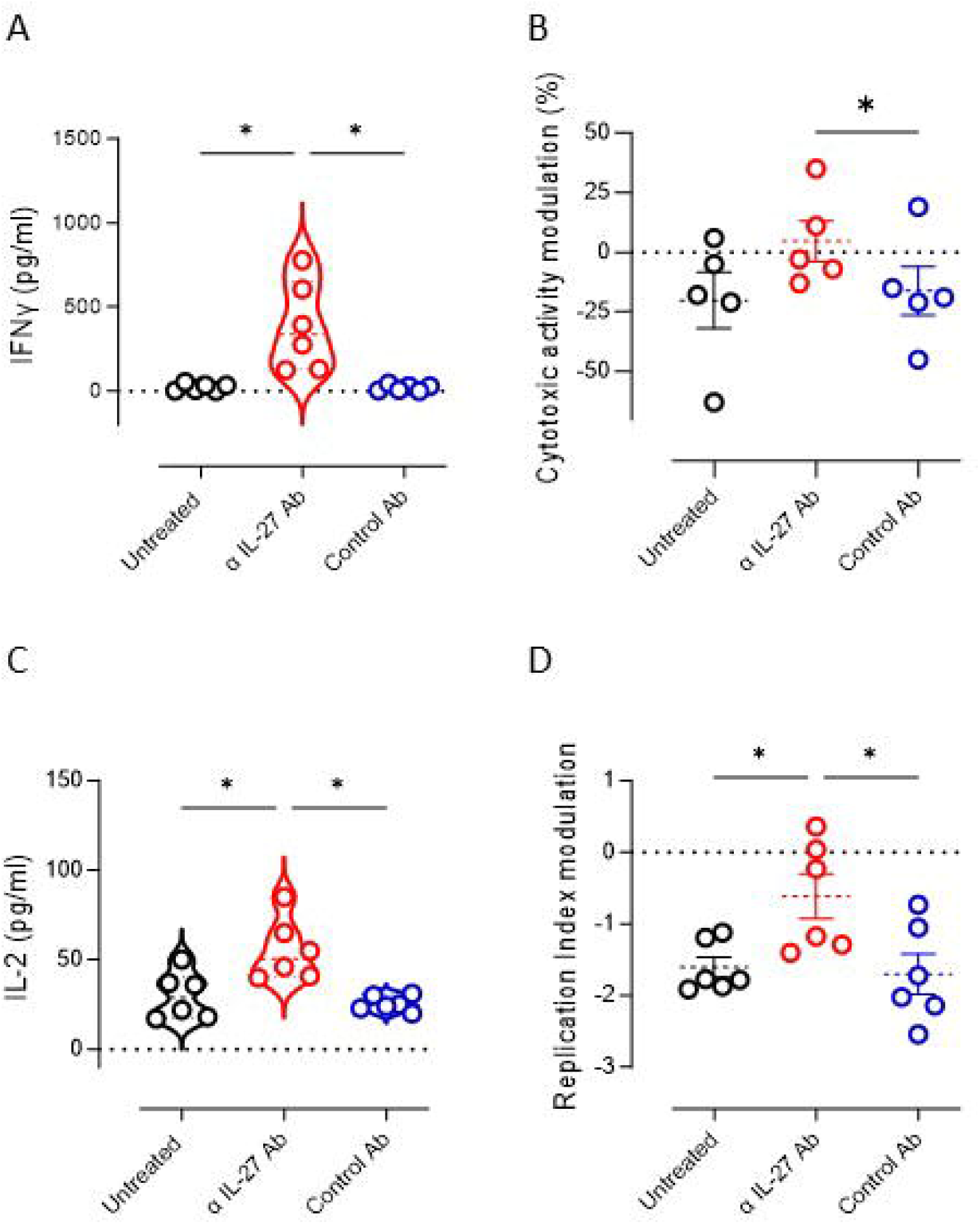
IL-27 neutralization restores macrophage support for T-cell activation and cytotoxicity. Macrophages (Untreated, αIL-27 Ab, Control Ab) were extensively washed and then LPS-activated for 24 h to generate M-CM; matched isotype IgG was included in all cultures. (A) IFN-γ secretion by activated CD8⁺ T cells cultured with 10% macrophage M-CM (ELISA). (B) Macrophage effects on the cytotoxicity of a MUC1-specific CD8⁺ T-cell clone after 72 h co-culture with tumor targets. Macrophages were differentiated with M-CSF (Untreated), M-CSF + αIL-27 Ab, or M-CSF + isotype control (Control Ab), washed, and then added to T cells and targets. Cytotoxicity is shown as the percent change in tumor-cell lysis relative to the no-macrophage control (set to 0%); negative values indicate suppression and positive values enhancement. (C) IL-2 secretion by activated CD4⁺ T cells cultured with 10% M-CM (ELISA). (D) Impact of 10% M-CM on CD4⁺ T-cell replication index after 6 days (CFDA-SE dilution by flow cytometry). ELISA in ng/mL. Each dot = an independent experiment; mean ± SEM. Statistics: one-way ANOVA with Tukey’s post-hoc. Significance: p < 0.05 (*p<0.05).

Next, we investigated the effect of macrophages on CD8⁺ T cell cytotoxicity using an in vitro tumor cell killing assay. In the absence of macrophages, the MUC1-specific CD8⁺ T cells efficiently lysed the MUC1⁺ tumor cells (41). Adding Miso macrophages to the co-culture reduced the overall cytotoxicity of the T cells (approximately a 10% decrease in tumor cell killing compared to no macrophage; Figure 6B). In contrast, adding Mα27 macrophages did not impair CD8⁺ T cell cytotoxicity; the lysis levels remained comparable to the no-macrophage condition (Figure 6B). Moreover, in direct comparison, activated Mα27 macrophages were significantly less suppressive than activated Miso macrophages, resulting in ∼19% higher T-cell cytotoxicity in the presence of Mα27 (Figure 6B). These results indicate that IL-27-neutralized macrophages, unlike control macrophages, do not inhibit CD8⁺ T cell tumor-killing capacity. All assays were performed under the same washed/isotype-controlled conditions described above.

CD4⁺ helper T cells are critical for sustaining CD8⁺ T cell responses, partly through IL-2 production. We therefore examined IL-2 secretion by CD4⁺ T cells in the presence of M-CM. After 48 h of anti-CD3/CD28 stimulation, CD4⁺ T cells secreted nearly twice as much IL-2 when cultured with supernatant from activated Mα27 macrophages compared to supernatant from Miso macrophages (Figure 6C). This suggests that factors from IL-27-neutralized macrophages are less inhibitory (or possibly more supportive) to CD4⁺ T cell activation. In line with this, we assessed CD4⁺ T cell proliferation over 6 days by CFDA-SE dilution. We found that M-CM from activated Mα27 macrophages caused a significantly smaller reduction in the CD4⁺ T cell replication index than M-CM from activated Miso or from untreated macrophages (Figure 6D). In other words, CD4⁺ T cells proliferated more robustly in the presence of factors from IL-27-neutralized macrophages. As above, these experiments were conducted after thorough macrophage washing and with matched isotype controls in all conditions.

These experiments demonstrate that IL-27 neutralization endows macrophages with a diminished capacity to suppress T cell responses. CD8⁺ T cells maintain higher IFN-γ production and cytotoxic function, and CD4⁺ T cells exhibit greater IL-2 secretion and proliferation, in the context of Mα27 versus Miso macrophages. This improved support for T cell activity is consistent with the reduced output of immunosuppressive mediators (e.g., IL-10, PGE₂) and increased output of supportive factors (e.g., GM-CSF) by IL-27-neutralized macrophages.

## Discussion

In this study, we identify intratumoral IL-27 as a regulator of an immunosuppressive network in the tumor microenvironment. Using the MC38 colon adenocarcinoma model, we show that local IL-27 neutralization delays tumor growth, prolongs survival, and enhances the efficacy of PD-L1 blockade. Mechanistically, intratumoral IL-27 blockade reduced CD39 expression across several tumor-infiltrating immune populations, including myeloid cells and CD8⁺ T cells, and decreased the CD39^high^ fraction among CD4⁺ T cells.

In parallel, our human macrophage data demonstrate that IL-27 neutralization during M-CSF-driven differentiation impairs the acquisition of an immunoregulatory phenotype. This is evidenced by the downregulation of markers such as CD39, CD73, CD155, IL-10, CCL22, and PGE₂/COX-2, alongside an upregulation of pro-inflammatory molecules including CD80, CD40, CD48, and GM-CSF. Importantly, this polarization shift is accompanied by a metabolic reprogramming toward glycolysis, reinforcing a pro-inflammatory metabolic state. Functionally, IL-27-neutralized macrophages were less suppressive toward CD4⁺ and CD8⁺ T cells, allowing improved IL-2 production, proliferation, IFN-γ secretion, and cytotoxic activity. Together, these data support a model in which local IL-27 sustains a suppressive myeloid and adenosinergic program that limits anti-tumor immunity and reduces responsiveness to immune checkpoint blockade.

The dual role of IL-27 in cancer has long been recognized but remains mechanistically difficult to reconcile (42). IL-27 can promote type 1 immunity, CD8⁺ T cell survival, cytotoxic differentiation, and anti-tumor responses in several experimental settings. Conversely, IL-27 can also induce immunoregulatory pathways, including IL-10 production, inhibitory receptor expression, PD-L1 expression, and ENTPD1/CD39 upregulation (1,2,7–10). Our data support the idea that these apparently divergent outcomes are not mutually exclusive, but instead reflect differences in anatomical compartment, cellular target, timing, dose, and mode of IL-27 exposure.

This distinction is particularly important in light of the recent study by Bréart et al., which demonstrated a potent CTL-supportive role for IL-27 in anti-tumor immunity (5). In that study, IL-27 expression correlated with CTL signatures in human and murine tumors, IL-27 directly supported tumor-specific CD8⁺ T cell persistence and effector function, and therapeutic IL-27 receptor agonism using inducible IL-27 expression or half-life-extended IL-27 induced tumor regression and synergized with PD-L1 blockade. High IL-27 expression was also associated with favorable outcome in patients treated with anti-PD-1/PD-L1 therapy. These findings convincingly establish that IL-27 can act as a CTL-supportive cytokine when the signal reaches the relevant CD8⁺ T cell compartment with sufficient magnitude and duration.

Our findings do not contradict this CTL-supportive function. Rather, they reveal a distinct compartment-specific role of IL-27 within the established tumor bed. Bréart et al. mainly highlight the consequences of systemic IL-27 pathway manipulation and therapeutic agonism, including effects on tumor-specific CTLs and draining lymph node responses (5). In contrast, our approach was designed to neutralize endogenous IL-27 locally within the tumor microenvironment. This is a critical experimental difference. Systemic IL-27 blockade may simultaneously remove IL-27 signals required for CTL priming, persistence, and effector programming in lymphoid and tumor compartments. By contrast, intratumoral neutralization is expected to preferentially disrupt the local IL-27-dependent suppressive circuitry operating within the tumor niche, while minimizing interference with systemic or lymph node IL-27 functions.

This framework helps reconcile why IL-27 agonism, systemic IL-27 gene delivery, systemic IL-27 blockade, and intratumoral IL-27 neutralization can produce different outcomes (42). Strong systemic IL-27 exposure can impose a dominant CTL program, characterized by enhanced cytotoxicity, improved CD8/Treg ratio, and in some models Treg depletion (5,6). By contrast, chronic endogenous IL-27 production within an established tumor may participate in a negative feedback loop that limits inflammation and reinforces suppressive pathways (42). In this local setting, IL-27 may act on macrophages, dendritic cells, Tregs, and dysfunctional T cells to sustain CD39 expression, adenosine generation, IL-10 production, checkpoint ligand expression, and reduced T cell activity. Thus, the biological output of IL-27 is likely determined not only by the presence of the cytokine, but by the balance between its CTL-intrinsic stimulatory effects and its myeloid/Treg-associated regulatory effects.

The CD39/CD73/adenosine pathway provides a mechanistic bridge between these observations. CD39 hydrolyzes extracellular ATP and ADP into AMP, which can then be converted by CD73 into adenosine, a potent inhibitor of anti-tumor immunity (16–18). CD39 also marks highly suppressive regulatory T cells and terminally exhausted tumor-infiltrating CD8⁺ T cells (19–21). Previous work demonstrated that IL-27 induces ENTPD1/CD39 in dendritic cells and contributes to T cell suppression (11). IL-27-driven CD39 expression also confers protumorigenic activity to regulatory T cells (12). In macrophages, our previous work showed that IL-27 contributes to CD39 upregulation during M-CSF-driven differentiation and promotes acquisition of an immunoregulatory phenotype (13). The present study extends these observations in vivo by showing that local IL-27 neutralization reduces CD39 expression across multiple tumor-infiltrating immune subsets.

Our human macrophage data further support a model in which IL-27 acts upstream of a broader suppressive macrophage program. IL-27 neutralization reduced CD39 and CD73, suggesting impaired capacity to generate adenosine. It also decreased CD155, a ligand involved in TIGIT-dependent immune inhibition, and reduced IL-10, CCL22, and PGE₂/COX-2, all of which are associated with suppressive or tumor-promoting macrophage functions. Conversely, IL-27 neutralization increased CD80, CD40, CD48, and GM-CSF, consistent with a macrophage state more permissive to T cell activation. These phenotypic changes were accompanied by metabolic reprogramming, with increased mitochondrial respiration, ATP-linked respiration, and lactate production. Although macrophage metabolism cannot be reduced to a simple M1/M2 dichotomy, these changes indicate a more activated and bioenergetically engaged state. Functionally, IL-27-neutralized macrophages were less able to suppress T cell activation and cytotoxicity, linking the phenotypic and metabolic changes to an improved effector immune response.

The improvement of PD-L1 blockade after intratumorally IL-27 neutralization can therefore be explained by the coordinated weakening of several suppressive pathways. First, reduced CD39 expression on myeloid cells and T cells may limit adenosine production and decrease adenosine-mediated inhibition of effector lymphocytes. Second, reduced CD39 on CD8⁺ T cells may reflect attenuation of terminal exhaustion-associated features. Third, decreased CD39 intensity within the CD4⁺ compartment may indicate reduced suppressive potential among CD39^high^ regulatory-like cells, although FoxP3⁺ Tregs were not directly quantified in our in vivo panel. Fourth, IL-27-neutralized macrophages produced fewer soluble suppressive mediators, including IL-10 and PGE₂, and were less inhibitory toward T cells in vitro. Together, these effects likely create a tumor microenvironment more permissive to PD-L1 blockade, even if IL-27 neutralization may also reduce PD-L1 expression on macrophages, as shown previously (13). In this context, the net therapeutic benefit may result from reducing macrophage-driven and adenosine-dependent suppression rather than from increasing PD-L1 target density.

Our results also have implications for how IL-27-targeted therapies should be conceptualized. The data do not support a simplistic view in which IL-27 is either uniformly anti-tumoral or uniformly protumoral. Instead, they suggest that therapeutic outcome may depend on whether the intervention amplifies IL-27 signaling in CTLs or blocks IL-27-dependent suppression within the tumor microenvironment. Systemic IL-27 agonism may be beneficial when the goal is to directly program tumor-specific CTLs, increase cytotoxicity, and improve the CD8/Treg balance (5,6). Conversely, local IL-27 neutralization may be beneficial in tumors where endogenous IL-27 primarily sustains macrophage, Treg, and exhausted T cell suppressive circuits through CD39 and related pathways. Therefore, IL-27 expression alone may not be sufficient as a biomarker. A more informative strategy may be to assess IL27 together with EBI3, ENTPD1/CD39, NT5E/CD73, IL10, myeloid infiltration, Treg enrichment, and CTL signatures to distinguish IL-27-associated inflamed tumors from IL-27-dependent suppressive niches. Future studies should directly compare intratumorally and systemic IL-27 neutralization side-by-side in the same tumor model. Such experiments would clarify whether systemic blockade impairs CTL priming or maintenance while local blockade preferentially disrupts suppressive myeloid and adenosinergic circuits

In conclusion, our study identifies local endogenous IL-27 as a driver of CD39-associated immune suppression in the tumor microenvironment. By neutralizing IL-27 directly within the tumor, we reduce a coordinated suppressive network involving macrophages, CD39^high^ myeloid cells, exhausted-like CD8⁺ T cells, and CD39^high^ CD4⁺ T cells, thereby improving anti-tumor immunity and sensitizing tumors to PD-L1 blockade. These findings complement, rather than oppose, recent studies showing that systemic or therapeutic IL-27 agonism can support CTL-mediated tumor control. Together, the literature and our data support a compartmental model in which IL-27 can either promote anti-tumor CTL immunity or reinforce local immunosuppressive feedback, depending on dose, kinetics, anatomical localization, and cellular target. This model provides a rationale for developing IL-27-based strategies that either agonize CTL-supportive IL-27 signaling or locally block IL-27-dependent suppressive circuits according to the immune architecture of the tumor.

## Declarations

### Ethics approval

Animal experiments in this study have been approved by the ethics committee on which Institute Infinity depend (APAFIS#28910-2021010716077736 v2).

In this study, we used human cells isolated from blood for which we have agreement ANG-2003-2 and 25 21PLER2021-010.

### Availability of data and material

All data relevant to the study are included in the article or uploaded as supplementary information. Data are available upon reasonable request.

### Competing interest

None

### Funding

This work was performed in the context of the LabEX IGO program, supported by the National Research Agency through the investment for the future program ANR-11-LABX-0016-01, and by grants from the Ligue Contre le Cancer (Conseil Scientifique Inter Regional Grand-Ouest, CSIRGO and Occitanie) of the integrated research program 3I-Impact, supported by the University of Angers and the University Hospital of Angers. Supported by institutional grants from Inserm, CNRS, the University of Angers, the University of Toulouse and the University Hospital of Angers.

### Authors’ contributions

Conception and design JT, YD, QG

Experiment realization QG, LB, PP, TB, SdA

Data analysis and interpretation QG, YD, CB, JT

Funding acquisition JT, YD

Supervision JT, YD

Writing original draft JT, QG

JT is responsible for the overall content as guarantor

## Acknowledgements

We thank all members of mice core facilities (UMS006, ANEXPLO, Inserm, Toulouse) for their support and technical assistance. We thank Dominique Rozet for the daily help on the administrative and financial management of our team. The authors are grateful to the CRCI2NA technical platforms for the support and to the We-Met platform for the Seahorse analysis. Flow cytometry experiments were performed at the Univ Toulouse, INSERM, CNRS, Infinity core facility connected to Toulouse Réseau Imagerie network, member of the France-BioImaging national infrastructure supported by the French National Research Agency (ANR-24-INBS-0005 FBI BIOGEN). We thank all team members for their availability and help.

## List of Abbreviations

A2AR: Adenosine A2A Receptor
Ab: Antibody
AMP: Adenosine Monophosphate
ANOVA: Analysis of Variance
ATP: Adenosine Triphosphate
BRET: Bioluminescence Resonance Energy Transfer
CCL: Chemokine Ligand
CCR: C-C Chemokine Receptor
CD: Cluster of Differentiation
CFDA-SE: Carboxyfluorescein Diacetate Succinimidyl Ester
COX: Cyclooxygenase
CTLA-4: Cytotoxic T-Lymphocyte Antigen-4
DC: Dendritic Cells
DNase: Deoxyribonuclease
ECAR: Extracellular Acidification Rate
ECL: Enhanced chemiluminescence
ELISA: Enzyme Linked Immunosorbent Assay
EP2/4: Prostaglandin E2 receptor 2/4
GAPDH: Glyceraldehyde 3-Phosphate Dehydrogenase
GM-CSF: Granulocyte Macrophage-Colony Stimulating Factor
gMFI: Geometric Mean of Fluorescence Intensity
ICI: Immune Checkpoint Inhibitors
IFNγ: Interferon Gamma
IL: Interleukine
LAG3: Lymphocyte-Activation Gene 3
LPS: Lipopolysaccharide
mAbs: Monoclonal Antibodies
M-CSF: Macrophage-Colony Stimulating Factor
M-CM: Macrophages-Conditioned Medium
MDSC: Myeloid-Derived Suppressor Cells
MFI: Median Fluorescence Intensity
mRNA: Messenger RNA
MUC1: Mucin-1
NK: Natural Killer
OCR: Oxygen Consumption Rate
OxPhos: Oxidative Phosphorylation
PBS: Phosphate-Buffered Saline
PCR: Polymerase Chain Reaction
PD-1: Programmed Death 1
PDGF: Platelet-Derived Growth Factor
PD-L1: Programmed cell Death-Ligand 1
PGE2: Prostaglandin E2
PHA: Phytohemagglutinin
PTGS2: Prostaglandin-endoperoxide Synthase 2
RFI: Relative Fluorescence Intensities
RNA: Ribonucleic Acid
RPMI: Roswell Park Memorial Institute
SEM: Standard Error of the Mean
TAM: Tumor-Associated Macrophages
TGFβ: Tumor Growth Factor β
Th1: T helper 1
TIGIT: T cell Immunoreceptor with Ig and ITIM domains
TIM3: T-cell Immunoglobulin and Mucin-domain containing 3
TME: Tumor Microenvironment
TNFα: Tumor necrosis factor alpha
Treg: T regulator cell
VEGF: Vascular Endothelial Growth Factor

## Supplementary Figure legends

**Supplementary Figure 1. Expression and secretion of IL-27 by human monocytes and macrophages.** mRNA expression of IL-27 subunits (IL-27p28, Ebi3) and receptor subunits (IL-27RA, gp130) in monocytes prior to differentiation, quantified by RT-qPCR and normalized to GAPDH (A). Induction of IL-27p28 and IL-27RA transcripts in monocytes after 5 days of differentiation with M-CSF, assessed by RT-qPCR and normalized to GAPDH (B). IL-27p28 concentration in culture supernatants of monocytes and macrophages, with or without LPS stimulation, measured by ELISA (C). Membrane-associated IL-27p28 in unstimulated monocytes, measured by ELISA (D). Membrane-associated IL-27p28 during macrophage differentiation (day 0 to day 5, M-CSF), measured by ELISA (E).Results are expressed as mean ± SEM of independent experiments.

**Supplementary Figure 2. CD39 expression on macrophages after treatment with immunomodulatory compounds.** Macrophages differentiated for 7 days with M-CSF were exposed to the following modulators: (A) the adenosine receptor antagonist caffeine, (B) the EP2 receptor antagonist AH6809, and (C) GM-CSF. Surface CD39 was quantified by flow cytometry. Results are expressed as relative fluorescence intensity (RFI), calculated as MFI_specific / MFI_isotype (isotype-matched control). Each dot represents an independent experiment; data are mean ± SEM. Statistics: one-way ANOVA with Tukey’s post-hoc test (two-tailed; α = 0.05). Significance: *p* < 0.05 *(*p<0.05, **p<0,01, ****p<0,0001)*.

**Supplementary Figure 3. IL-27 neutralization during macrophage differentiation modifies phenotype.** Macrophages were generated for 7 days with M-CSF (Untreated), M-CSF + **α**IL-27 antibody (**α**IL-27 Ab), or M-CSF + isotype-matched IgG (Control Ab). (A) *NT5E* (CD73) mRNA by RT-qPCR (normalized to GAPDH). (B) CCL2 secreted after 24 h LPS stimulation (ELISA), (C) CCL2 secreted by inflammatory macrophages (with and without activation step), (D) IL-10 secreted after 24 h LPS stimulation (ELISA). (E) *PTGS2* (COX-2) mRNA by RT-qPCR. (F) COX-2 protein by western blot (representative donor). (G) Paired comparison of COX-2 protein levels relative to Untreated. Each dot represents an independent experiment; data are mean ± SEM. Statistics: one-way ANOVA with Tukey’s post-hoc test (paired analysis used where donor-matched). Significance: *p* < 0.05 (*p<0.05, **p<0,01).

**Supplementary Figure 4. Macrophage conditioned medium inhibits IFN-γ secretion by CD8⁺ T cells.** In the indicated condition, LPS-activated M-CM of untreated macrophages (10% of total volume) was added to cultures of CD8^+^ T cells along with PHA at 10 µg/mL for 48 h. In Mock condition no conditioned medium was added. IFN-γ in culture supernatants (ELISA). Each dot represents an independent experiment; data are mean ± SEM. Statistics: one-way ANOVA with Tukey’s post-hoc test (two-tailed; α = 0.05). Significance: *p* < 0.05 (*).

**Supplementary Figure 5. Gating strategy for flow-cytometry analysis of the tumor microenvironment.** C57BL/6 mice bearing subcutaneous MC38 tumors were analyzed by flow cytometry 10 days after inoculation. The figure shows the gating sequence used to identify tumor-infiltrating macrophages, monocytes, neutrophils, dendritic cells, CD4⁺ T cells, and CD8⁺ T cells. A representative experiment is shown. Analysis was performed in FlowJo v10; instrument details are provided in Methods.

## Notes

### Competing Interest Statement

The authors have declared no competing interest.

### Summary of Updates

We are submitting a revised version of our manuscript because we have conducted a thorough reanalysis of our data. This process has led to updated conclusions that differ from those presented in the previous version. Additionally, the data presentation has been significantly revised to better reflect these new findings and improve clarity. We believe these changes are essential to ensure the accuracy and robustness of our study, and we appreciate the opportunity to share this updated version with the scientific community.

