## supplemental figures for "IL-27 neutralization promotes pro-inflammatory macrophage polarization and improve response to immune checkpoint blockade"

### Slide 1
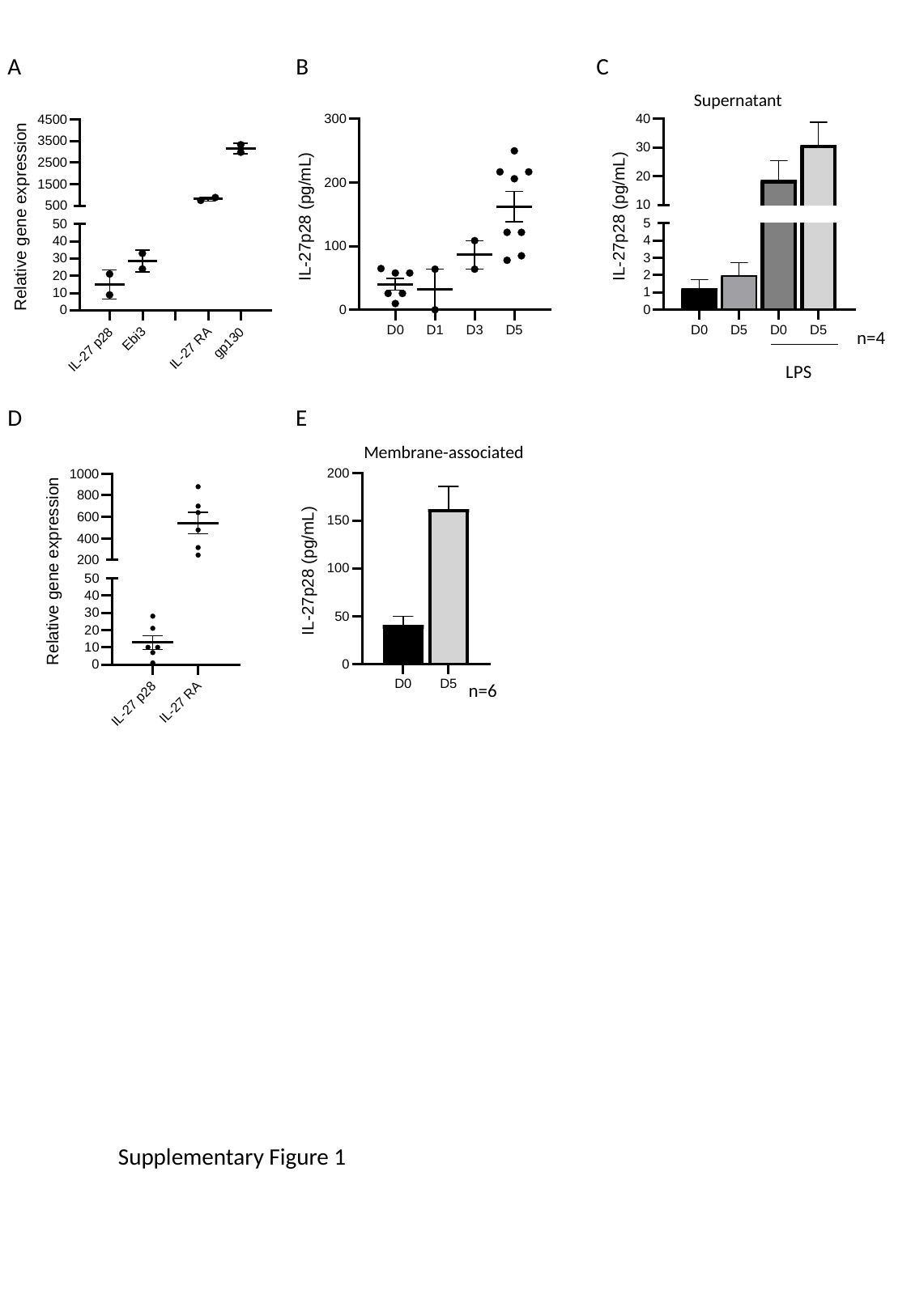

A
B
C
Supernatant
n=4
LPS
D
E
Membrane-associated
n=6
Supplementary Figure 1

### Slide 2
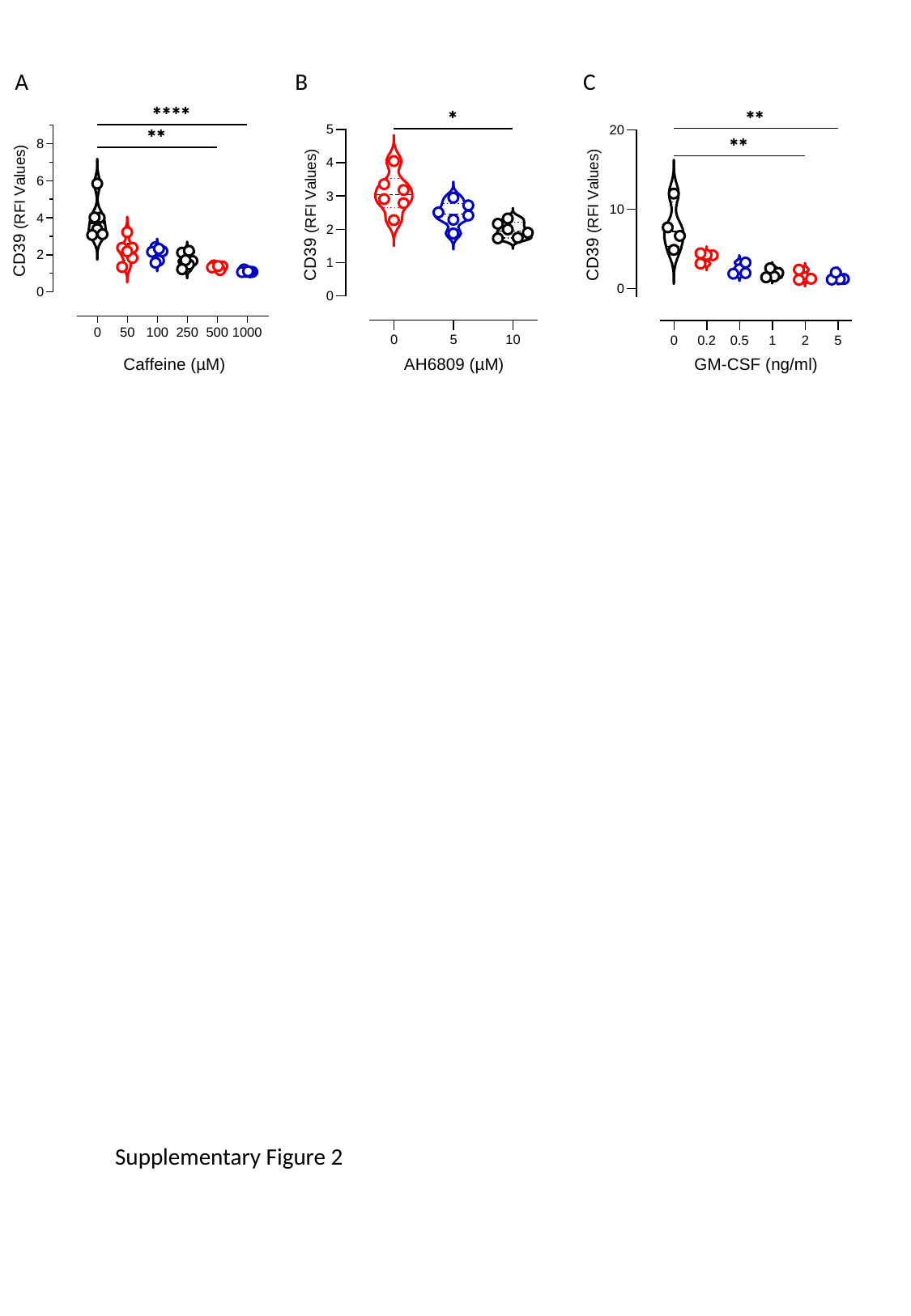

A
B
C
Supplementary Figure 2

### Slide 3
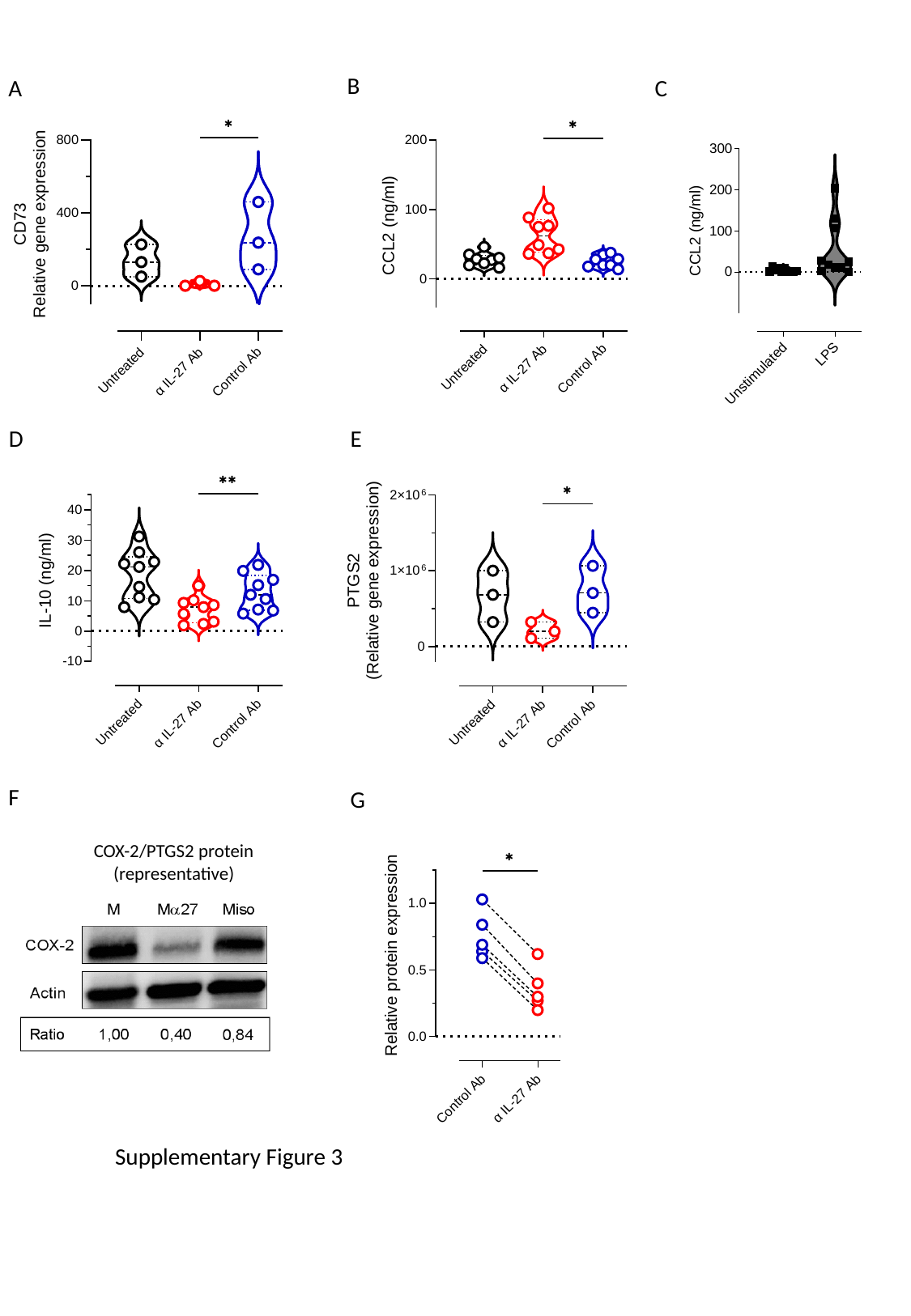

B
C
A
D
E
F
G
COX-2/PTGS2 protein (representative)
Supplementary Figure 3

### Slide 4
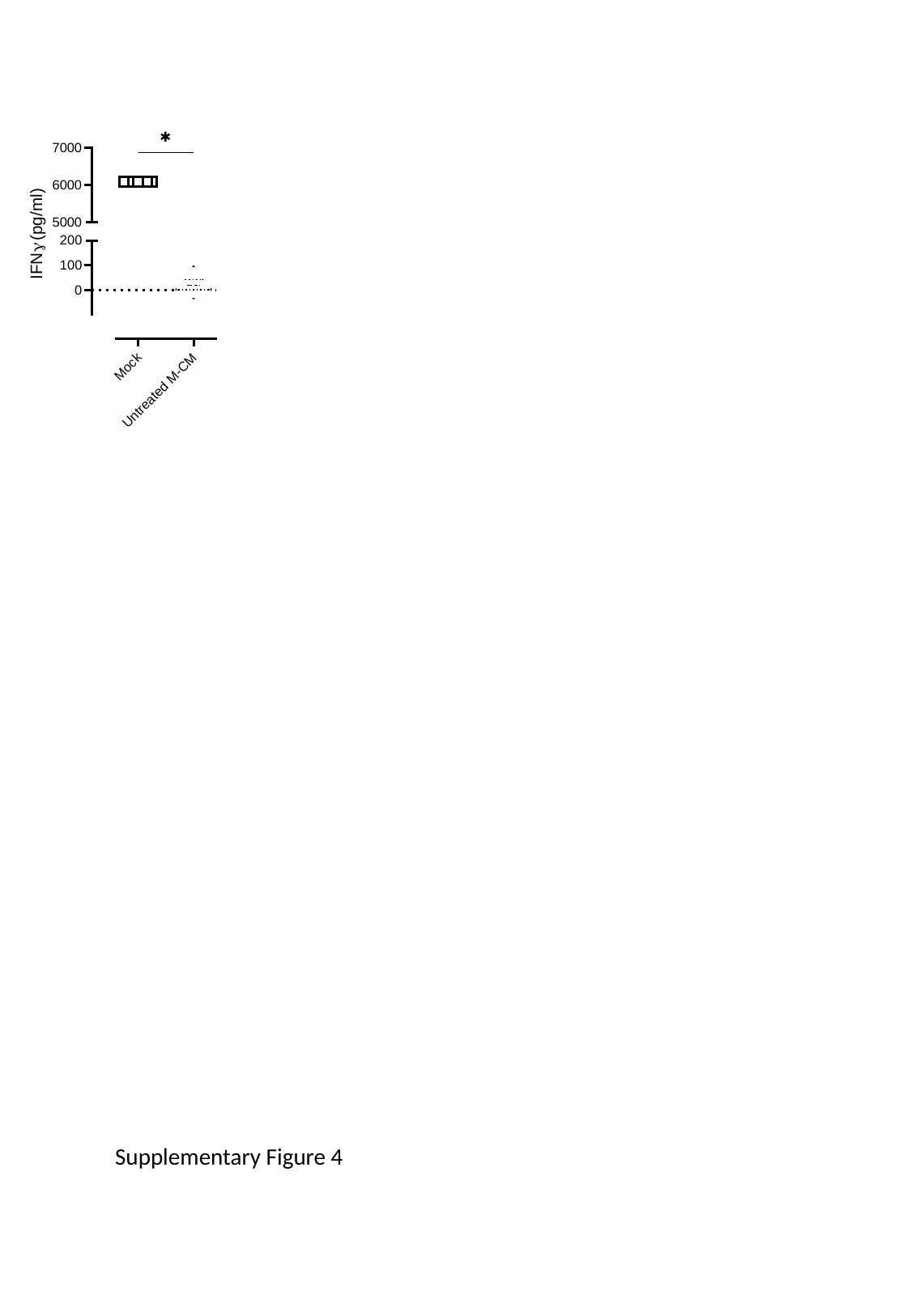

Supplementary Figure 4

### Slide 5
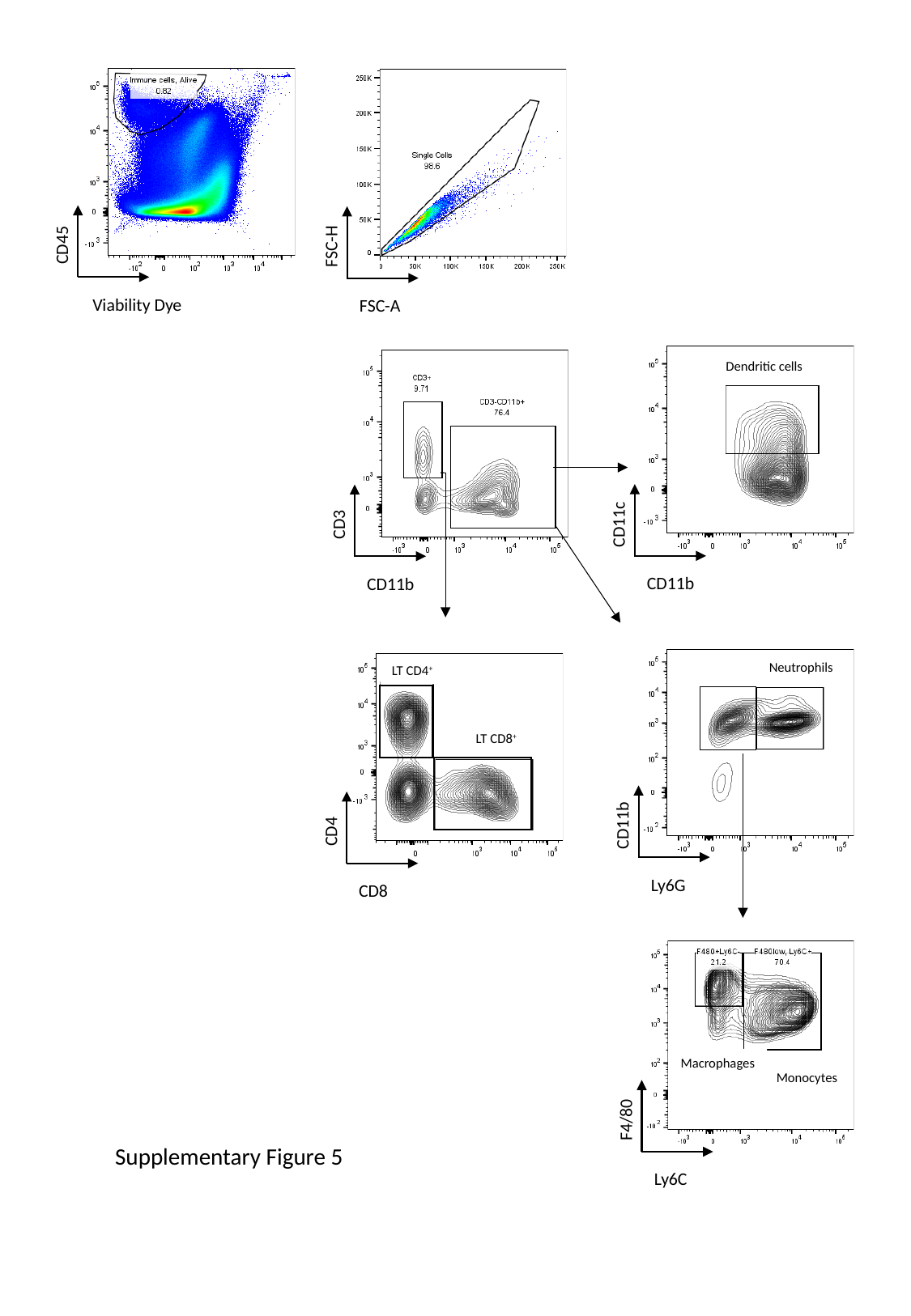

CD45
Viability Dye
FSC-H
FSC-A
CD11c
CD11b
CD3
CD11b
Dendritic cells
CD11b
Ly6G
CD4
CD8
Neutrophils
LT CD4^+^
LT CD8^+^
F4/80
Ly6C
Macrophages
Monocytes
Supplementary Figure 5

### Slide 6
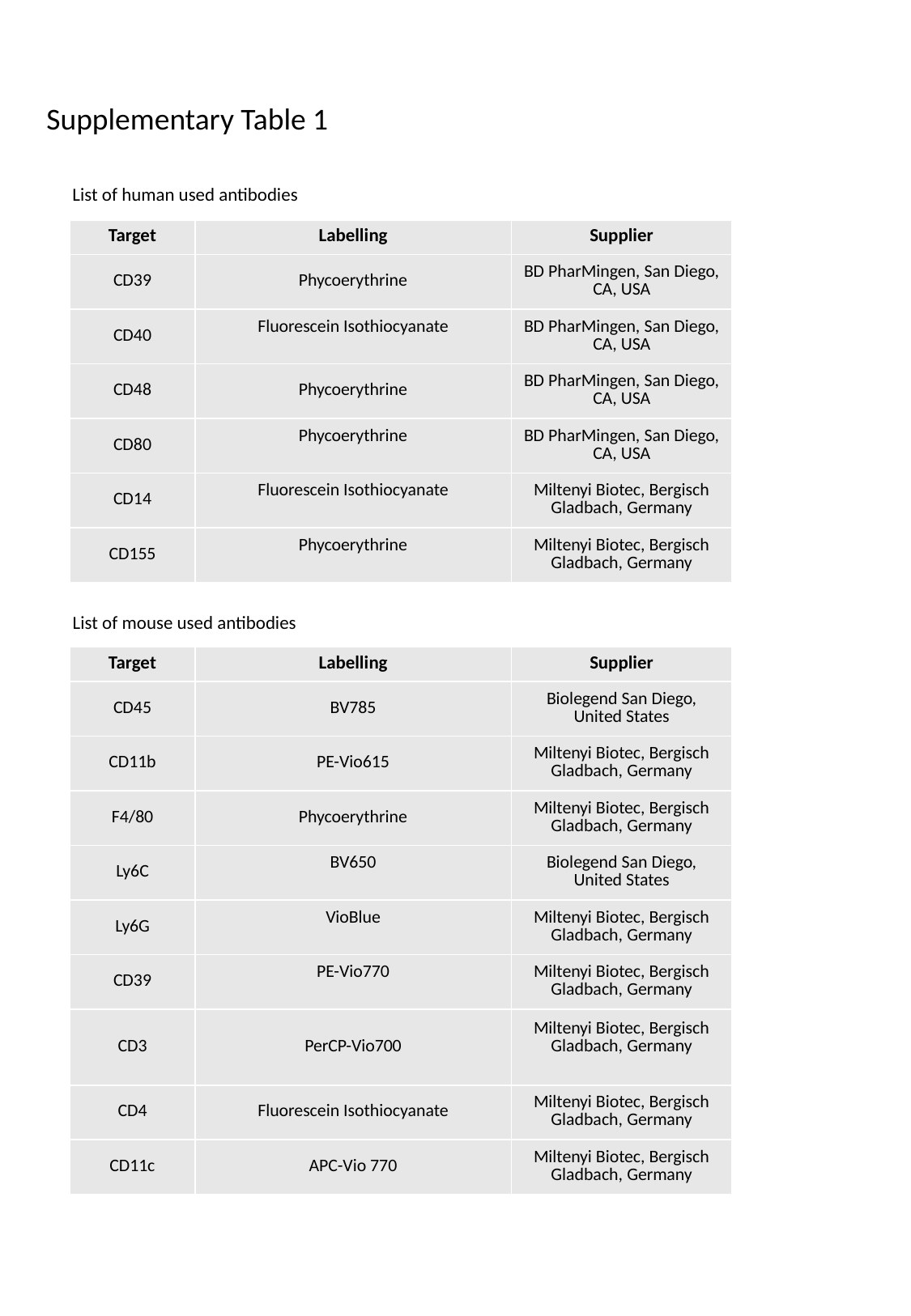

Supplementary Table 1
List of human used antibodies
| Target | Labelling | Supplier |
| --- | --- | --- |
| CD39 | Phycoerythrine | BD PharMingen, San Diego, CA, USA |
| CD40 | Fluorescein Isothiocyanate | BD PharMingen, San Diego, CA, USA |
| CD48 | Phycoerythrine | BD PharMingen, San Diego, CA, USA |
| CD80 | Phycoerythrine | BD PharMingen, San Diego, CA, USA |
| CD14 | Fluorescein Isothiocyanate | Miltenyi Biotec, Bergisch Gladbach, Germany |
| CD155 | Phycoerythrine | Miltenyi Biotec, Bergisch Gladbach, Germany |
List of mouse used antibodies
| Target | Labelling | Supplier |
| --- | --- | --- |
| CD45 | BV785 | Biolegend San Diego, United States |
| CD11b | PE-Vio615 | Miltenyi Biotec, Bergisch Gladbach, Germany |
| F4/80 | Phycoerythrine | Miltenyi Biotec, Bergisch Gladbach, Germany |
| Ly6C | BV650 | Biolegend San Diego, United States |
| Ly6G | VioBlue | Miltenyi Biotec, Bergisch Gladbach, Germany |
| CD39 | PE-Vio770 | Miltenyi Biotec, Bergisch Gladbach, Germany |
| CD3 | PerCP-Vio700 | Miltenyi Biotec, Bergisch Gladbach, Germany |
| CD4 | Fluorescein Isothiocyanate | Miltenyi Biotec, Bergisch Gladbach, Germany |
| CD11c | APC-Vio 770 | Miltenyi Biotec, Bergisch Gladbach, Germany |
